# Perinatal ampicillin administration modulates murine bile acid metabolism *in vivo* - an observational study

**DOI:** 10.1101/2024.01.31.578215

**Authors:** Sydney P Thomas, Fatemeh Askarian, Armin Kousha, Emi Suzuki, Chih Ming Tsai, George Liu, Victor Nizet, Pieter C Dorrestein, Shirley M. Tsunoda

## Abstract

Antibiotics are an indispensable tool of modern medicine, yet their impact extends beyond eliminating harmful bacteria to perturbing the commensal bacteria constituting the gut microbiome. This collateral damage is particularly significant in early life when the gut microbiome is still developing. In humans, antibiotic administration during infancy and childhood is associated with various long-term negative health outcomes. However, existing research has predominantly focused on the direct administration of antibiotics to infants, leaving uncertainties about whether indirect antibiotic exposure produces similar effects. Here, we use mouse models to investigate how three distinct routes of exposure to the commonly prescribed broad-spectrum antibiotic ampicillin influences parent and infant metabolism. These methods simulate major modes of both direct and indirect antibiotic exposure: intravenous antibiotic administration to the mother immediately before birth mimicking intrapartum antibiotic prophylaxis, antibiotic use by the mother during lactation, and direct administration to infants mimicking empiric antibiotic treatment for neonatal sepsis. Through untargeted metabolomics of fecal samples from mouse dams and infants, we identified one class of compounds, bile acids and related cholane steroids, as particularly sensitive to ampicillin treatment. Bile acids, produced by the host and extensively modified by the gut microbiome, serve as important mediators in the cross-talk between the microbiota and the host. Here, we detail the coordinated changes in bile acid metabolism in response to a commonly prescribed antibiotic, focusing on dams treated both pre- and postpartum. Additionally, we identify unique bile acids associated with weight gain in infant mice.

**Importance:** Antibiotics are widely used perinatally, administered to both parents and infants before, during, and after birth. While they can play a life-saving role, antibiotics also result in collateral damage to the beneficial microbes constituting the gut microbiome. These microbes have many important functions, particularly in the metabolism of small molecules in the body. One such group of molecules, bile acids, undergo extensive modifications by bacteria and may act as a “language” through which microbes communicate with the host. This observational study investigates the impact of the commonly prescribed antibiotic ampicillin on the metabolism of these molecules during childbirth. Our results indicate that ampicillin administration pre- or post-partum significantly alters the mother’s bile acid metabolism, but has a minimal influence on infant bile acid levels. However, in all cases, ampicillin administration significantly increased infant weight, even when the antibiotic was solely administered to the mother.

## Introduction

Worldwide, approximately 30% of infants and pregnant or lactating parents are prescribed antibiotics, although this percentage varies greatly based on geographical location.^1–7^ While these medications are an essential part of modern medicine, their overuse poses potential long-term health risks. These concerns are linked to disruptions in the gut microbiota — a complex and dynamic community of bacteria, fungi, archaea, protists, and viruses inhabiting the gastrointestinal tract. These microbes play crucial roles in metabolism and can be drivers or biomarkers of many diseases.^8,9^ While gut microbes have important functions throughout the lifespan, they have an outsized influence early in life.^10–12^ Development of the gut microbiota begins at birth and undergoes dynamic changes during infancy before stabilizing in adulthood.^13,14^ Antibiotics can influence the gut microbiota at any stage, but their administration during this crucial developmental window may lead to enduring changes in the gut microbial community.^15–19^ In humans, early life antibiotic use is associated with increased risk of conditions including obesity,^20–22^ Type 1 diabetes,^23^ inflammatory bowel disease,^24^ and asthma,^25^ among others.^26,27^ However, the vast majority of these studies have focused only on direct infant antibiotic administration,^14,21,23–25,28,29^ leaving gaps in understanding the potential effects of antibiotic use prepartum or during lactation.

Cross-talk between gut microbes and the host is often mediated by metabolites. Bile acids are a crucial part of this dialogue. The primary bile acids in mammals, cholic acid and chenodeoxycholic acid, are produced in the liver, but microbes transform them into a variety of secondary bile acids with diverse functions. Until recently, these secondary bile acids were thought to be the product of only four types of reactions: deconjugation of glycine or taurine, dehydroxylation, dehydrogenation, or epimerization of the bile acid core. However, many other bile acid conjugations have recently been observed, revealing a much greater diversity of bile acids than previously imagined.^30–34^ Thus, bile acids may represent a complex “language” used by microbes to communicate with the host. Until recently, this language remained hidden among the thousands of metabolic features in untargeted metabolomics whose chemical identities are unknown. To annotate these unknown molecules, experimental spectra must be matched to reference spectra. The recent development of a library of reference spectra for novel bile acids enables the annotation of molecules not previously identified. Given that many of these bile acids are likely produced by bacteria, they may be uniquely sensitive to microbiome perturbations following antibiotic exposure.

Here, we employed experimental mouse models to investigate three primary modes of exposure to the widely prescribed broad-spectrum antibiotic ampicillin (AMP) during early life: prepartum administration to the mother, postpartum administration to the mother during lactation, and direct infant administration (**Figure 1**). This parallel experimental setup enables a comparison of the impact of these different exposures on the metabolism of both the mother and infant. Our results uncover large alterations in bile acid metabolism in dams treated with antibiotics both pre- and postpartum, with less pronounced effects observed in pups. Nevertheless, in all three scenarios, pups exposed to antibiotics exhibited a significant increase in weight over the initial 21 days of life.

**Figure 1:**
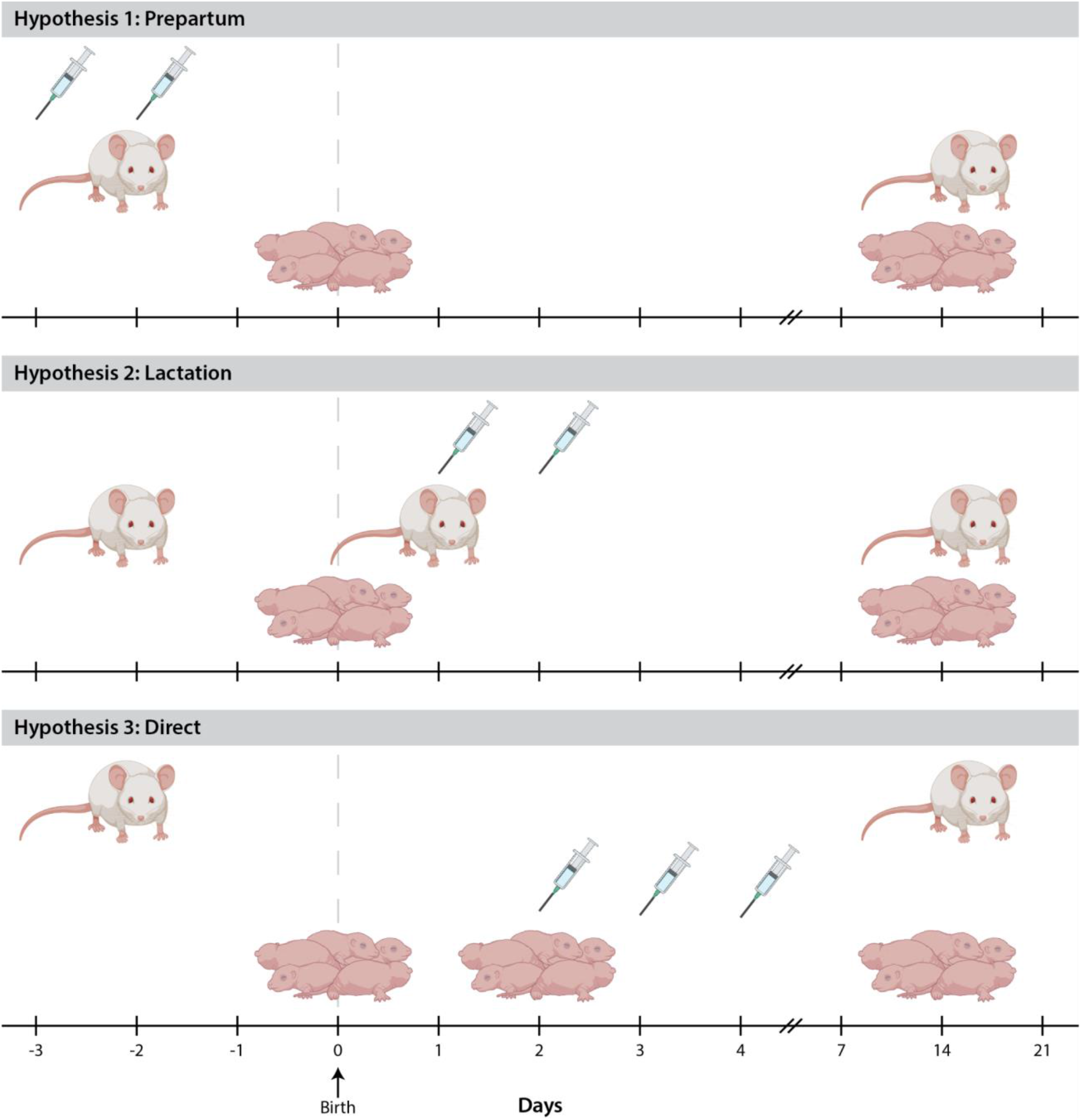
Experimental design. Mother and infant mice were treated intravenously with ampicillin to mimic three major modes of antibiotic administration in early life. Further details can be found in the Methods section. Mouse icons created using BioRender.com.

## Results

Our study encompassed three experimental arms (or Hypotheses), each representing a common method of antibiotic administration in early life (**Figure 1**). Hypothesis 1 involved mouse dams treated with ampicillin (AMP) prepartum. To reduce the risk of newborn group B streptococcal (GBS) infections, the American College of Obstetricians and Gynecologists recommends screening all pregnant women for GBS rectovaginal colonization between the 36th and 37th week of gestation, with the administration of intrapartum antibiotic prophylaxis (most commonly AMP) to all women whose cultures are positive unless a pre-labor cesarean section is performed in the setting of intact placental membranes.^35^ Hypothesis 2 treated mouse dams postpartum during lactation. Not infrequently, AMP is included as a component of multidrug regimens for treatment of serious postpartum maternal infections including surgical site infections, endometritis, septic thrombophlebitis, and urinary tract infection.^36^ Finally, Hypothesis 3 directly treated the infant mice. The World Health Organization recommends administering empiric AMP (and gentamicin) to neonates with documented risk factors for infection for 48 hours, continuing longer only in the case of positive cultures or clinical signs of sepsis.^37^

Fecal samples were collected longitudinally for both dams and pups, while serum samples were collected once at 21 days post-birth. TEMPoral TEnsor Decomposition (TEMPTED) for longitudinal sampling enabled the separation of fecal samples based on metabolic profile for dams treated with AMP both prepartum (Hypothesis 1) and postpartum (Hypothesis 2) while as expected no separation was observed for dams whose infants were directly administered AMP (Hypothesis 3) (**Figure 2a**).^38^ Out of the top 500 metabolic features distinguishing samples along Component 1, 64 were annotated using standard library matching, with exactly half of them (32) identified as bile acids (**Figure 2b**). Given the extensive modification of bile acids by the gut microbiome, we speculated that these molecules may be uniquely susceptible to antibiotic treatment. Rerunning TEMPTED with only the metabolic features matching bile acids provided comparable results to those obtained on the entire dataset (**Figure 2c**). Subject trajectories of dams treated with pre- or postpartum AMP revealed substantial changes in overall bile acid metabolism immediately after treatment, returning to baseline after 1-2 weeks (**Figure 2d**). This alteration involved groups of bile acids whose abundance increased or decreased directly after antibiotic administration, gradually reverting to pre-administration levels (**Figure S1A**). A subset of these bile acids included known primary and secondary bile acids, confirmed through retention time matching to pure standards (**Figure S2, Figure S3, Table S1**). Specifically, dams treated with AMP pre- or postpartum showed decreases in chenodeoxycholic acid and increases in taurocholic acid and glycolic acid (**Figure S2A**). Antibiotic administration had no significant impact on levels of these primary/secondary bile acids in pups (**Figure S2B**). We also identified several putative amino acid-conjugated bile acids, some of which were initially discovered in mice (**Figure S4**).^30^ Notably, Glu, Gly, Val, Leu/Ile, Phe, Tyr-conjugated bile acids were sensitive to antibiotic treatment, especially in dams treated with prepartum antibiotics (**Figure S4A**). The levels of many of these amino acid conjugates decreased immediately following antibiotic treatment before returning to pre-treatment levels. We only detected Val and Phe conjugates in pups.

**Figure 2:**
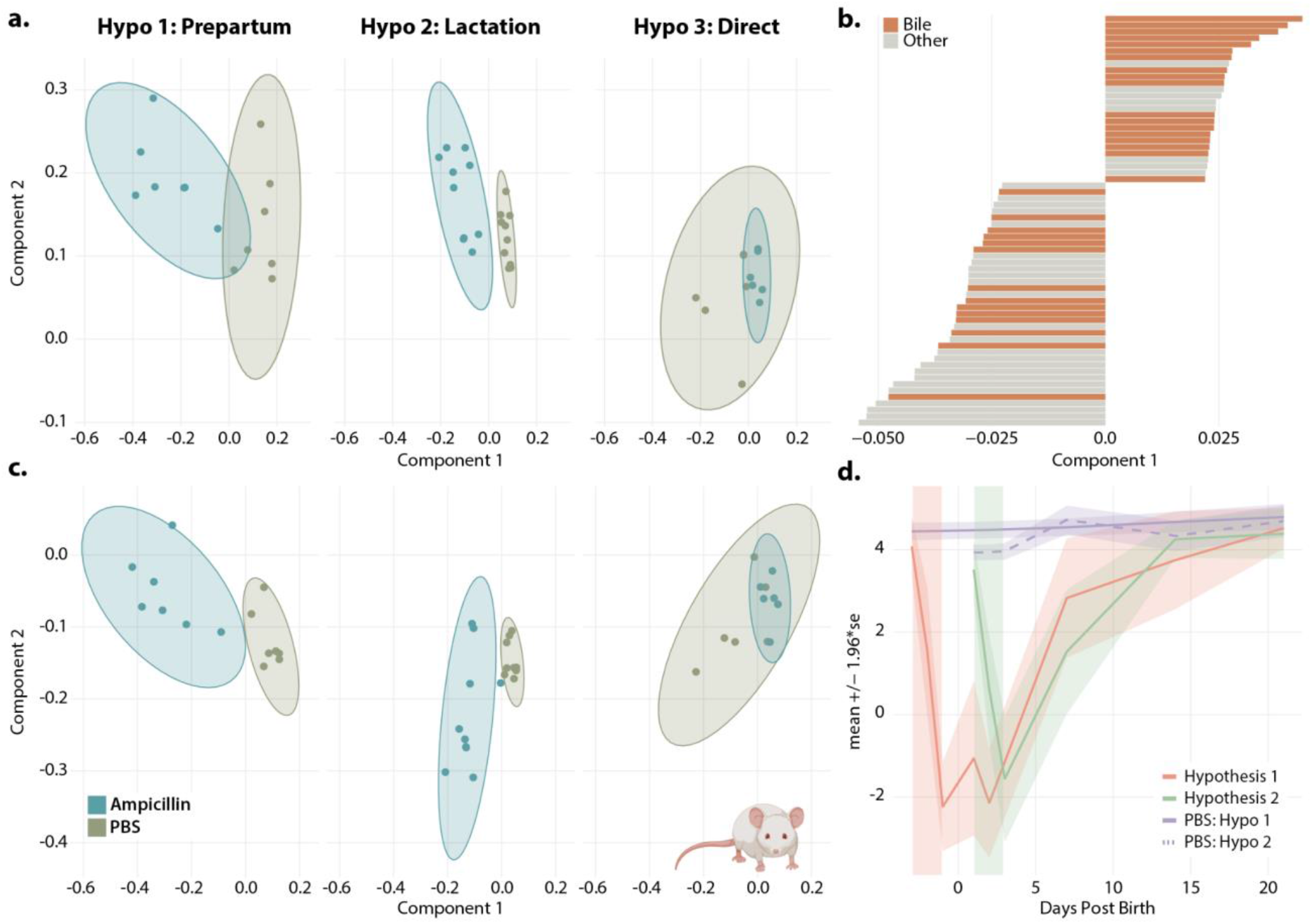
TEMPoral TEnsor Decomposition (TEMPTED) shows longitudinal differences in metabolism of mouse dams treated with ampicillin prepartum and postpartum. **a**. TEMPTED principal components using all metabolic features from untargeted metabolomics. **b**. Metabolic features with spectral library matches that drive separation along Component 1. Metabolites annotated as bile acids are shown in orange. **c**. Principal components using only metabolic features with library matches to bile acids **d**. Subject trajectories of dams treated with prepartum antibiotics (Hypo 1), postpartum antibiotics (Hypo 2), and controls (PBS). Colored blocks indicate the timing of ampicillin treatment.

In total, 354 metabolites were annotated as bile acids in all three experiments. Of these bile acids the majority (60%) were detected in the feces of both dams and pups, while the remaining 38% were identified in both feces and serum (**Figure 3A**). As many of these molecules were recently discovered, the chemical identity of the bile acid conjugate is often unknown. Nevertheless, bile acids can be classified based on the number of hydroxylations on the bile acid core. This classification revealed that di- and trihydroxylated bile acids are the dominant forms in both parent and infant mice, which aligns with the hydroxylation states of the major primary bile acids in mice (**Figure 3B**).^39^ In general, bile acids were more frequently detected in parent samples, likely attributed to the substantial changes in bile acid metabolism during the initial weeks of infant development (**Figure 3C-D**). These alterations are evident when comparing average bile acid abundances over time in the infant mice (**Figure S4B**). While none of the bile acids were statistically significant in the infant mice when comparing AMP and PBS treatment, a majority of bile acids exhibited significant differences at 21 days compared to earlier time points (**Figure S5B, S5D**).

**Figure 3:**
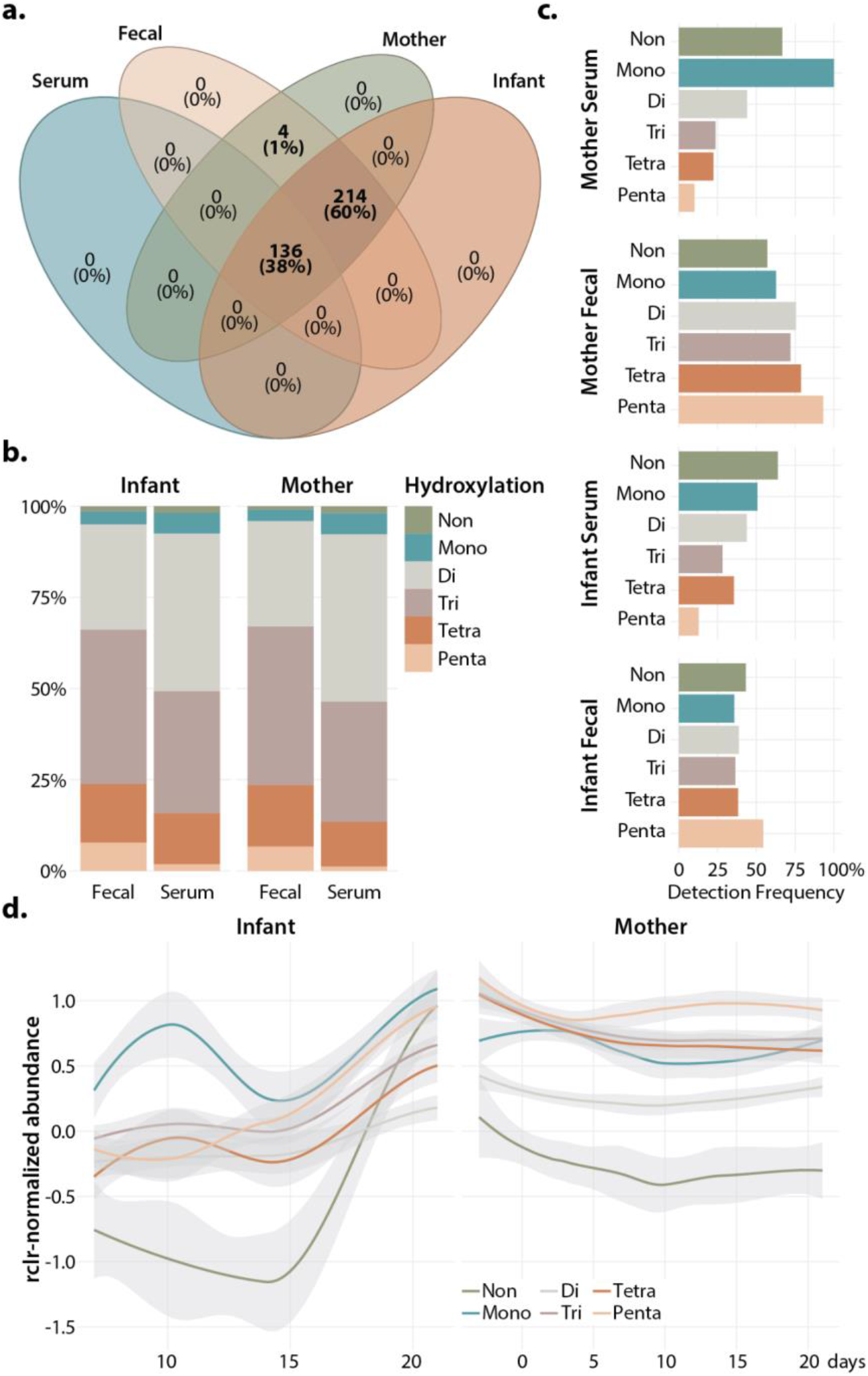
Overview of bile acid metabolism in mouse mothers and infants **a**. Number of bile acids observed in parent/infant serum and fecal samples **b**. Average abundance of bile acids with different levels of hydroxylation in parent/infant fecal and serum samples. **c**. Average detection frequency of bile acids with different levels of hydroxylations. **d**. Typical changes in hydroxylation levels during development. Average abundance of different levels of bile acid hydroxylation in control samples over time.

Although we measured over 350 bile acids in our experiments, not all molecules were detected in every sample. To understand how AMP administration influences the diversity of bile acids, we examined the number of bile acids detected in AMP-treated and control mice over time. AMP treatment significantly reduced the number of bile acids in dams treated pre- or postpartum but it had no effect on total bile acid abundance across any treatment (**Figure 4A**). Once again, infant mice did not exhibit significant differences in bile acid diversity or abundance following any route of AMP exposure (**Figure 4B**). This is consistent with the lack of significant differences in bile acid abundance when comparing AMP and PBS treatments (**Figure S5D**) and with the absence of separation between AMP-treated infants and controls using principal components analysis (**Figure S6**). However, infant mice did display significantly higher weights at Day 21 after all three forms of AMP exposure (**Figure 5A**). This effect was sex-specific depending on the type of antibiotic exposure. However, in both sexes and regardless of antibiotic exposure, we identified two bile acids that were significantly correlated to infant weight gain (**Figure 5B**). One of these bile acids matched to taurocholic acid sulfate, while the other matched an unknown conjugate with a mass difference of 55.018.

**Figure 4:**
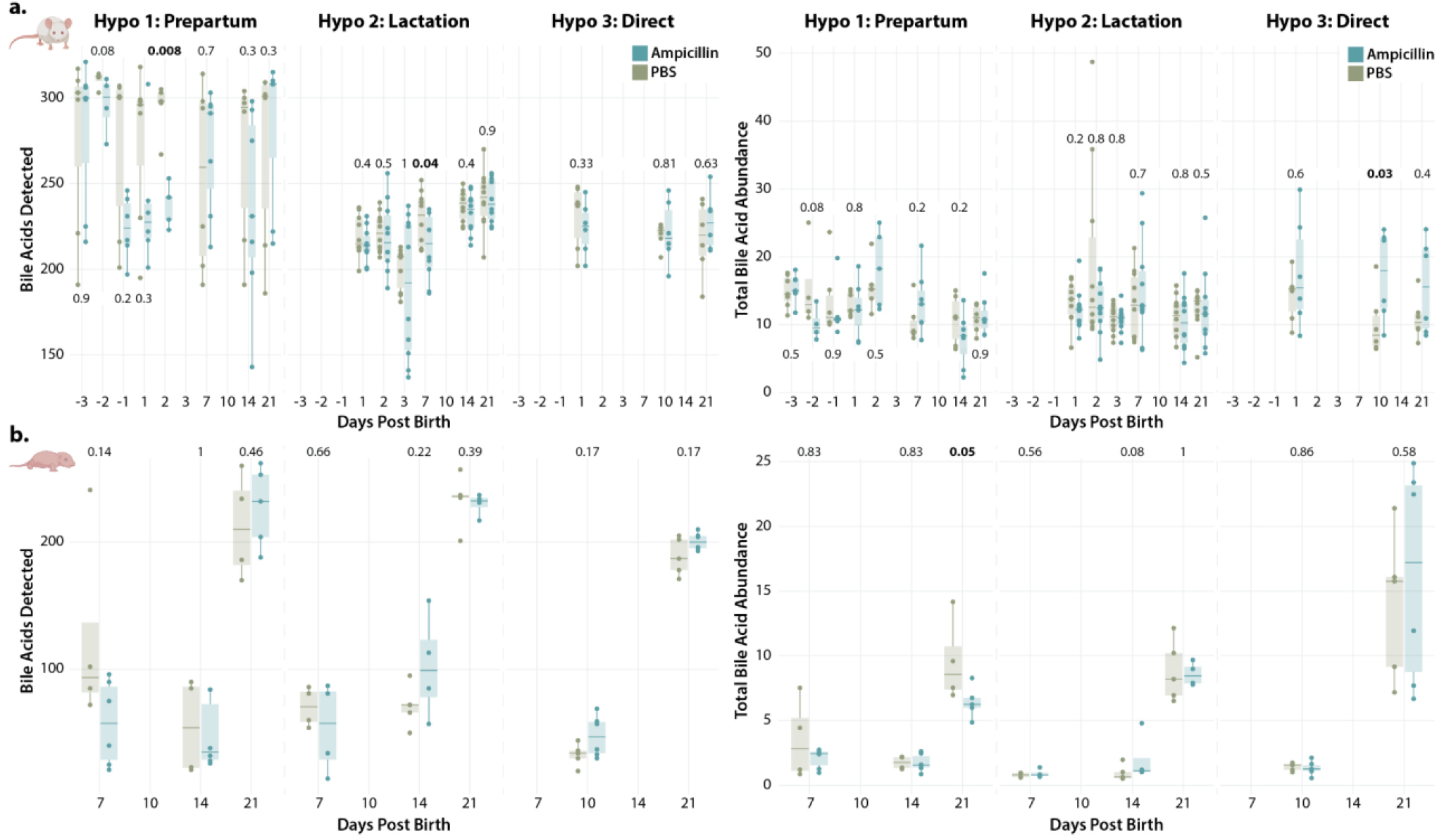
Diversity of bile acids, but not overall abundance, changes with ampicillin administration in parent mice. **a**. Number of bile acids detected (left) and total bile acid abundance (right) in parent mice. **b**. Number of bile acids detected (left) and total bile acid abundance (right) in infant mice. Total bile acid abundance calculated using total ion current (TIC)-normalized values.

**Figure 5:**
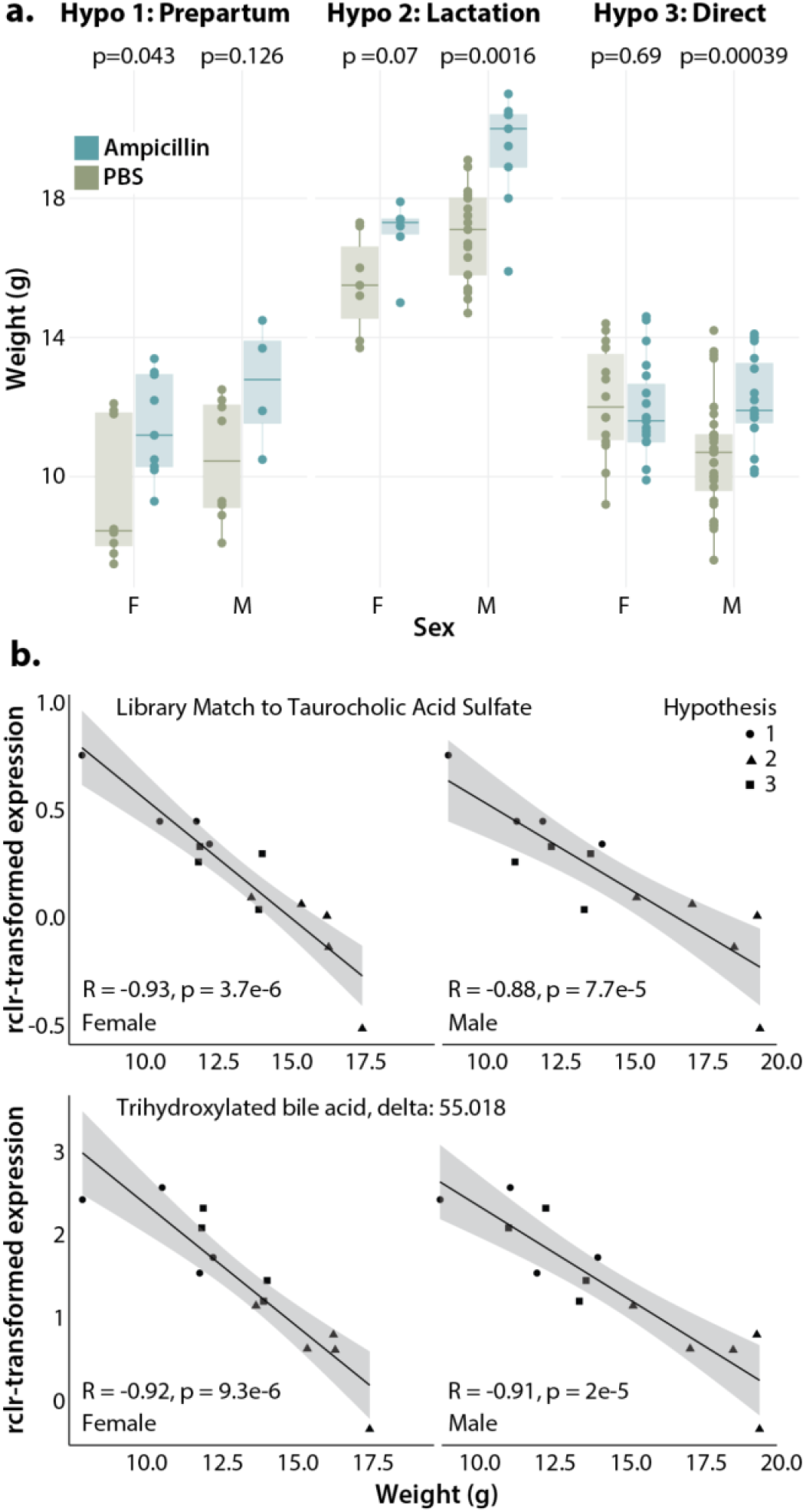
Infant weight increases after antibiotic treatment. **a**. Infant weights at 21 days with three methods of antibiotic administration. **b**. Putative bile acids with significant correlation to infant weights. Point shape indicates experimental group. Correlation calculated using spearman with BH correction.

## Discussion

Our results reveal a large and coordinated metabolic response to AMP in dams treated pre- or postpartum. Although these changes span several classes of metabolites, bile acids appear particularly sensitive to antibiotic administration. In fact, 146 of the 354 bile acids we detected (41%) exhibited significant differences in AMP-treated dams. These changes were remarkably consistent, as short asynchronous time-series analysis showed no significant differences in bile acid trajectories in dams treated with AMP pre- or postpartum (**Table S1**). Some of the bile acids we measured are well known, such as taurocholic acid and glycolic acid. These two bile acids increased significantly after antibiotic treatment, while levels of other candidate amino acid-conjugated bile acids decreased (**Figure S2 and S4**). In mice, the increase of taurocholic acid in response to antibiotic treatment has been previously shown to promote colonization of the fungal pathogen *Candida albicans*.^40–42^ In humans, taurocholic acid can alter blood glucose and has been associated with both liver cirrhosis and alcohol-associated liver disease.^43–45^ Both glycolic and taurocholic acid have been implicated in the progression of intrahepatic cholestasis of pregnancy, a severe complication affecting between 0.1-2% of pregnant women.^46–48^ While the increases in taurocholic and glycolic acid in this study were temporary, the effects of these antibiotic-induced changes on liver function merit further investigation. In addition to taurocholic and glycolic acid, the majority of bile acids that responded to antibiotic treatment were novel molecules whose biological functions have not yet been determined. However, since bile acids can act as signaling molecules to distal organs, these changes may have systemic effects beyond perturbations to the gut microbiome.^49^

Neither longitudinal nor standard principal components analysis separated out infant mice treated with AMP using either the entire untargeted metabolome or just those metabolites that matched bile acids (**Figure S6**). However, the significant increase in weight following AMP exposure suggests that AMP does alter the infant metabolome. This may be due to subtle shifts in metabolism that may not pass a significance threshold, but could also be due to technical difficulties in collecting fecal samples from infant mice. Due to the size of fecal pellets, we were unable to reliably collect any infant feces before 7 days of age, and unable to consistently collect pellets before 14 days of age. Thus, we could not capture changes in infant metabolism in the crucial early postnatal days directly after antibiotic administration in which we observed the largest differences in the dams. In addition, our data indicates that bile acid metabolism is highly dynamic during normal infant development. Thus, changes due to development may have obscured changes induced by AMP administration. In the future, collecting cecal contents from pups in the days immediately post antibiotic administration should provide greater clarity on what metabolic changes occur after AMP administration. However, we were able to identify two bile acids (taurocholic acid sulfate and a second unknown species) that were negatively correlated to weight regardless of antibiotic treatment. Bile acids have been implicated in both weight gain and weight loss, and play an important role in the complex metabolic response to diet and other environmental factors.^50–52^ The identification of novel bile acids associated with weight may help to untangle the mechanism behind weight gain after antibiotic exposure.^53^

## Materials and Methods

### Murine model of ampicillin administration

Mice were split into three cohorts to replicate the three major modes of antibiotic exposure in early life (see **Figure 1**).

#### Prepartum

Timed-pregnant C57BL/6 dams (12-14 weeks) were treated intravenously via retro-orbital injection with 150 mg/kg ampicillin (AMP) (n=5) or PBS (n=7) on days 16 and 17 of gestation (mice gave birth on day 19 of gestation).

#### Postpartum/Lactation

Timed-pregnant C57BL/6 dams (12-14 weeks) were treated intravenously via retro-orbital injection with 150 mg/kg AMP (n=9) or PBS (n=9) on days 1 and 2 post-delivery.

#### Direct

C57BL/6 pups were treated intraperitoneally with 150 mg/kg AMP (n=38 infants from 5 timed-pregnant dams) or PBS (n=46 from 6 timed-pregnant dams) on days 2, 3, and 4 post birth.

Each experiment was run twice to reduce batch effects and reach at least n=5 samples for each timepoint of each experimental group. All experiments were run concurrently with a PBS-injected control group. Mice were housed in filter-top cages with access to a chow diet and water under conditions of regulated ambient temperature (20-22 °C) and relative humidity (30-70%), with a 12 h light/12h dark cycle over the course of the experiment. Murine sera were collected from freshly drawn blood via submandibular bleeding at day 21 post birth. The serum was collected in BD Microtainer serum separator tubes (10 min, 10000 × *g*, 4 °C) by centrifugation. Fecal pellets were collected at multiple time intervals as indicated in Figure 1. Serum and feces were stored at -80 °C immediately after collection.

### Ethics declarations

Mouse experiments conformed to all ethical guidelines for animal research and were carried out in accordance with the rules and regulations of the Institutional Animal Care and Use Committee, which was approved by the UC San Diego IRB protocol S00227M.

### Metabolomics

#### Extraction

Fecal samples were extracted in 10 uL cold 50:50 MeOH:H2O per 1 mg fecal pellet. Pellets were agitated in a tissue lyzer at 25 Hz for 5 minutes and then incubated at 4C for 30 minutes. Finally, samples were centrifuged at max speed for 3 minutes and supernatant was transferred to 96 well plates.

For serum, 100 uL MeOH was added to 25 uL serum and vortexed before centrifuging at max speed for 3 minutes. Then 75 uL of the supernatant was added to 75 uL cold H2O in a 96 well plate. All solvents were purchased from Thermo Fisher and were LC-MS grade.

#### LC-MS/MS

Untargeted metabolomics was performed using data dependent acquisition (DDA) with an UltiMate 3000 liquid chromatography system (Thermo Scientific) coupled to a QExactive Orbitrap (Thermo Scientific) mass spectrometer. Samples were separated using a Kinetex C18 column (Phenomenex) and run in the positive mode. Metabolite separation was performed with a linear gradient of mobile phase A (water 0.1% formic acid (v/v)) and phase B (acetonitrile 0.1% formic acid (v/v)). A representative linear gradient was run at 0-1 min isocratic at 5% B, 1-7 min to 98% B, 7-7.5 min isocratic at 98% B, 7.5-8 min to 5% B, and 8-10 min at 5% B with a flow rate of 0.5 mL/min. All solvents used were LC-MS grade. Full scan MS spectra were acquired at 35,000 resolution with an automated gain control (AGC) target of 5e5, maximum ion injection time of 100 ms, and a scan range of 100-1500 m/z. MS/MS spectra were collected using the same resolution and AGC target, and fragmented the top 5 most abundant ions per cycle with a 3.0 m/z isolation window, stepped normalized collision energies of 20, 30, and 40%, and a dynamic exclusion window of 10 s.

To accurately identify known primary and secondary bile acids, we ran purified standards of the following bile acids: chenodeoxycholic acid, cholic acid, glycochenodeoxycholic acid, glycocholic acid, glycohyodeoxycholic acid, hyodeoxycholic acid, taurocholic acid, taurochenodeoxycholic acid, taurodeoxycholic acid, and taurohyodeoxycholic acid. Retention time matches to these standards are available in **Figure S3**. Bile acids with matched retention time standards are referred to by their chemical name; otherwise, metabolic features are labeled as a candidate bile acid. Since many of these molecules were only recently discovered, the chemical identity of the bile acid conjugate is often unknown. In this case, the conjugate is represented by its delta mass, or the difference in mass between the conjugated bile acid and its unconjugated core.

Feature finding and library matching was performed using GNPS and MZmine3.^54,55^ Metabolomics datasets are publicly available in the MassIVE database (http://massive.ucsd.edu) under MSV000092652, MSV000089558, and MSV000093192.

### Statistical analysis

Statistical analysis was performed in R. Scripts detailing the analysis are available on GitHub: https://github.com/Sydney-Thomas/Antibiotics_Affect_BA_Metabolism.

## Data availability

All data required to evaluate the paper’s conclusions are present in the paper and supplementary information. Metabolomics data are publicly available through Massive (http://massive.ucsd.edu) under the following dataset identifiers:

Hypothesis 1 (prepartum dams and infants): MSV000089558 and MSV000093192

Hypothesis 2 (postpartum/lactation dams and infants): MSV000092652 and MSV000093192

Hypothesis 3 (direct exposure infants and dams): MSV000093192

MZmine3 outputs are included in the Massive datasets and GNPS FBMN results can be accessed by the following link: https://gnps.ucsd.edu/ProteoSAFe/status.jsp?task=41dfdc8bc23342ee96347d6691483d4b.

Code and data tables used for statistical analysis and plotting are available on Github: https://github.com/Sydney-Thomas/Antibiotics_Affect_BA_Metabolism.

## Acknowledgements

This work was supported by NIH grants T32HD087978 and P50HD106463.

## Supplementary Information

The supplementary information contains Supplementary Figures S1-S6 and expanded metabolomics data associated with this manuscript (Table S1).

### Description of Table S1

VIPs_Parent_TEMPTED_all: Annotations and principal components for all metabolic features in parent fecal samples from Hypothesis 1 (prepartum) and Hypothesis 2 (lactation). Metabolites labeled TRUE in Component 1 or Component 2 are among the top 7.5% most important variables for each Principal Component.

VIPs_Parent_TEMPTED_bile: Annotations and principal components for all bile acids in parent fecal samples from Hypothesis 1 (prepartum) and Hypothesis 2 (lactation). Metabolites labeled TRUE in Component 1 or Component 2 are among the top 7.5% most important variables for each Principal Component.

SantaR_Hypo1_v_Hypo2: Annotations and p values for all bile acids in parent fecal samples from Hypothesis 1 (prepartum) and Hypothesis 2 (lactation). P values calculated using Short Asynchronous Time-Series Analysis (SantaR).

Parent_ANOVA_AMP: Annotations and p values comparing AMP treatment and PBS controls at each time point for all bile acids in parent fecal samples from Hypothesis 1 (prepartum), Hypothesis 2 (lactation), and Hypothesis 3 (direct). P values calculated using Tukey’s HSD with BH correction.

Infant_ANOVA_AMP: Annotations and p values comparing AMP treatment and PBS controls at each time point for all bile acids in infant fecal samples from Hypothesis 1 (prepartum), Hypothesis 2 (lactation), and Hypothesis 3 (direct). P values calculated using Tukey’s HSD with BH correction.

Parent_ANOVA_Days: Annotations and p values comparing bile acid levels at different days post birth in parent fecal samples from Hypothesis 1 (prepartum), Hypothesis 2 (lactation), and Hypothesis 3 (direct). P values calculated using Tukey’s HSD with BH correction.

Parent_ANOVA_Days: Annotations and p values comparing bile acid levels at different days post birth in infant fecal samples from Hypothesis 1 (prepartum), Hypothesis 2 (lactation), and Hypothesis 3 (direct). P values calculated using Tukey’s HSD with BH correction.

**Figure S1:**
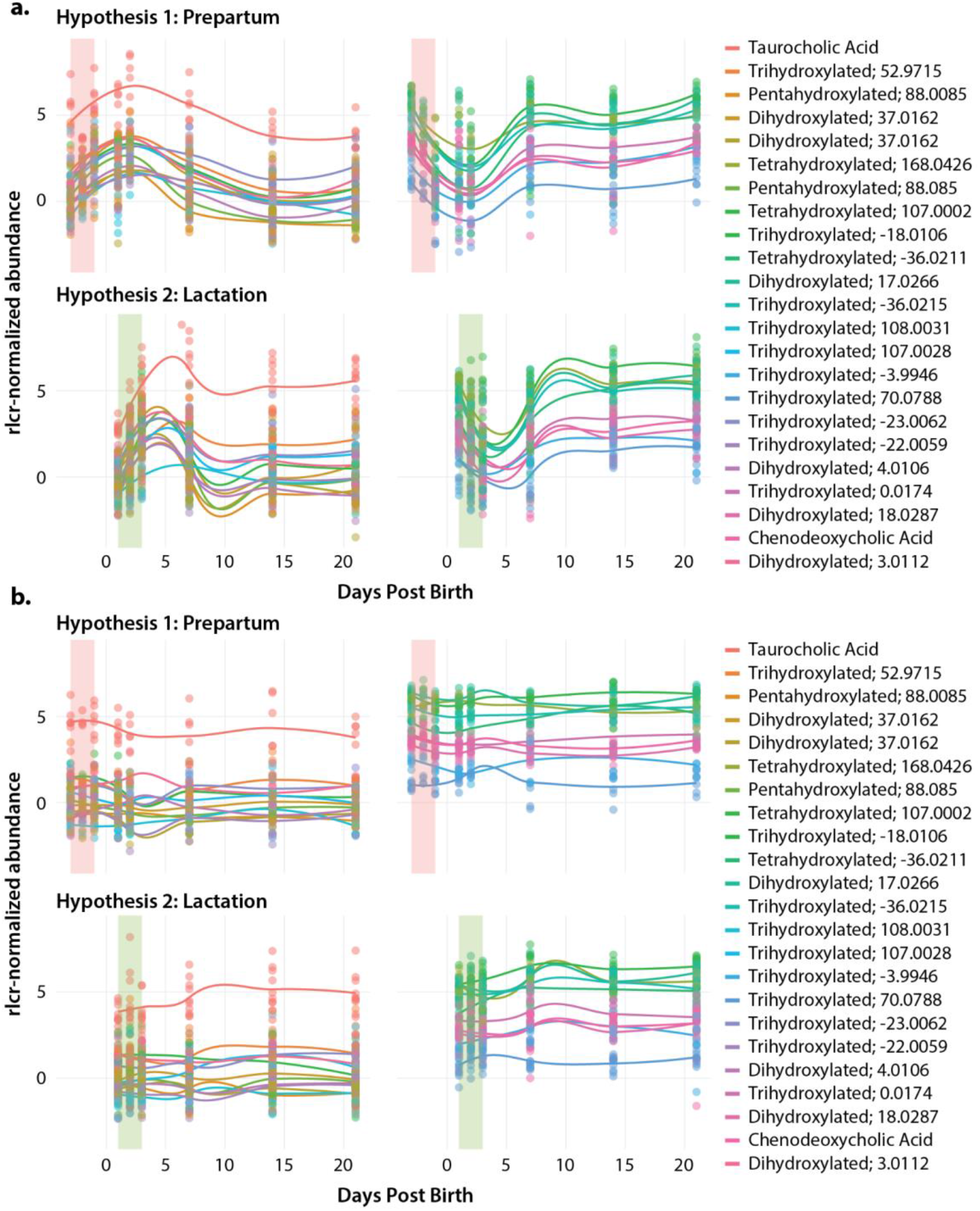
Trajectory of top 7.5% of bile acids driving separation along Component 1 in mouse dams **a**. treated with ampicillin or **b**. PBS controls (see Figure 2B). Lines indicate average abundance across all samples in a group, points indicate abundance in each individual. Colored bar indicates timing of antibiotic administration. Bile acids matched to purified retention time standards are labeled by their chemical name, otherwise, the conjugate is represented by hydroxylation and delta mass, or the difference in mass between the conjugated bile acid and its unconjugated core.

**Figure S2:**
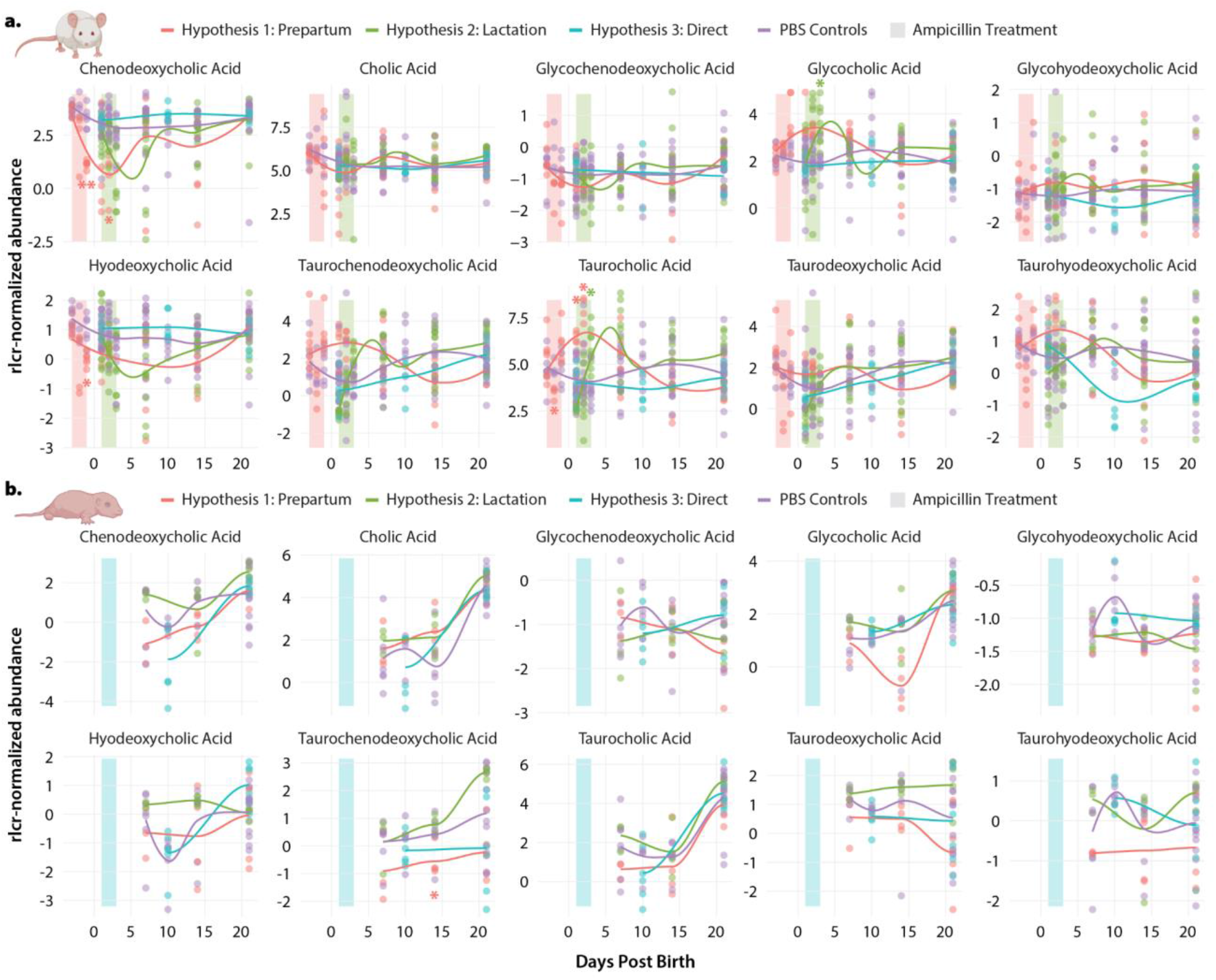
Abundance of primary and secondary bile acids confirmed using retention time matching in **a**. parent and **b**. infant mice feces. Lines indicate average abundance across the entire sample group, points indicate abundance of individual samples. Abundance is shown as robust centered log-transformed (rlcr-transformed) values. Significance calculated using Tukey’s HSD with BH correction. p.adj < 0.05 = *, p.adj < 0.01 = **, p.adj < 0.001 = ***.

**Figure S3:**
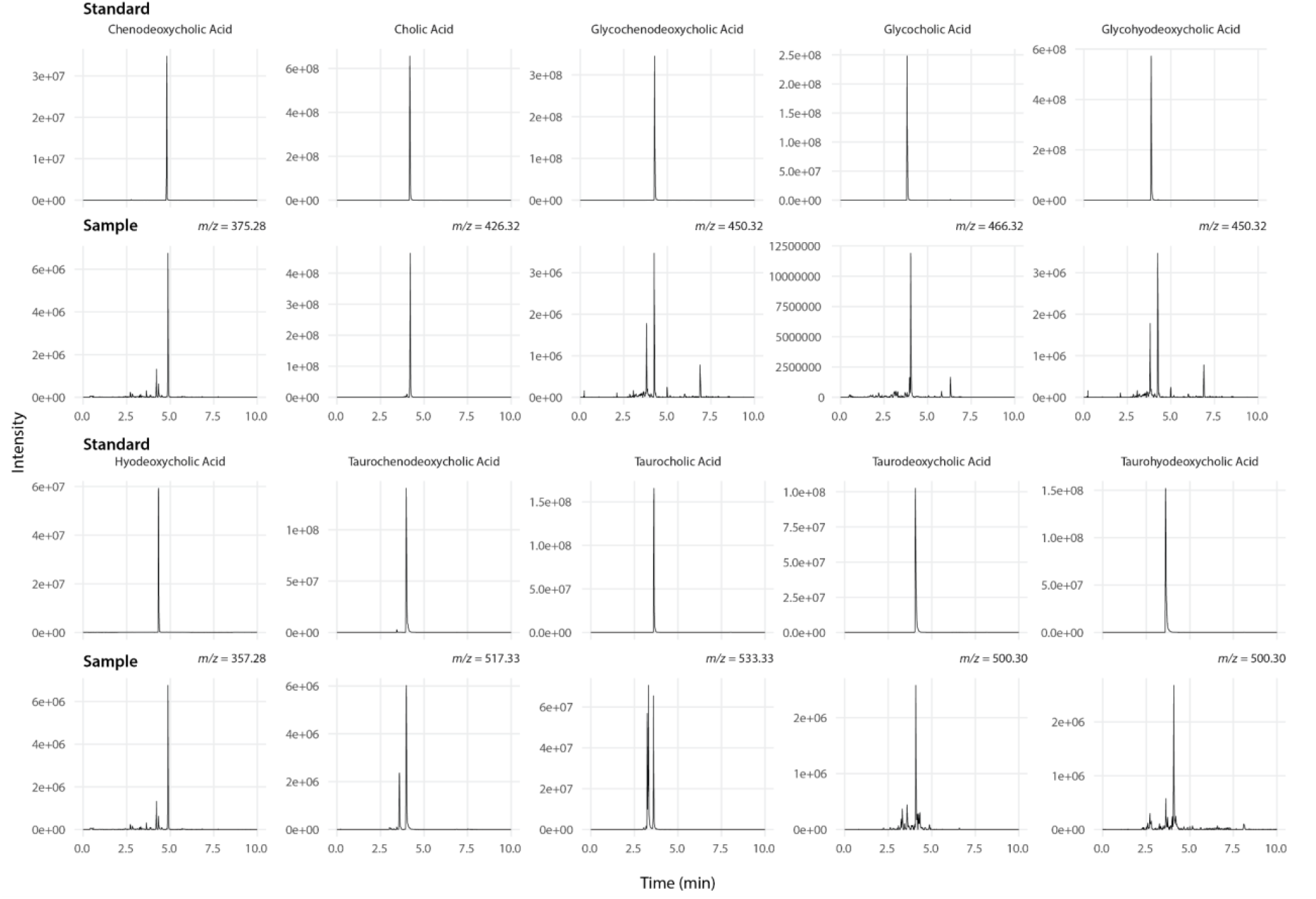
Retention time matches to commercial bile acid standards.

**Figure S4:**
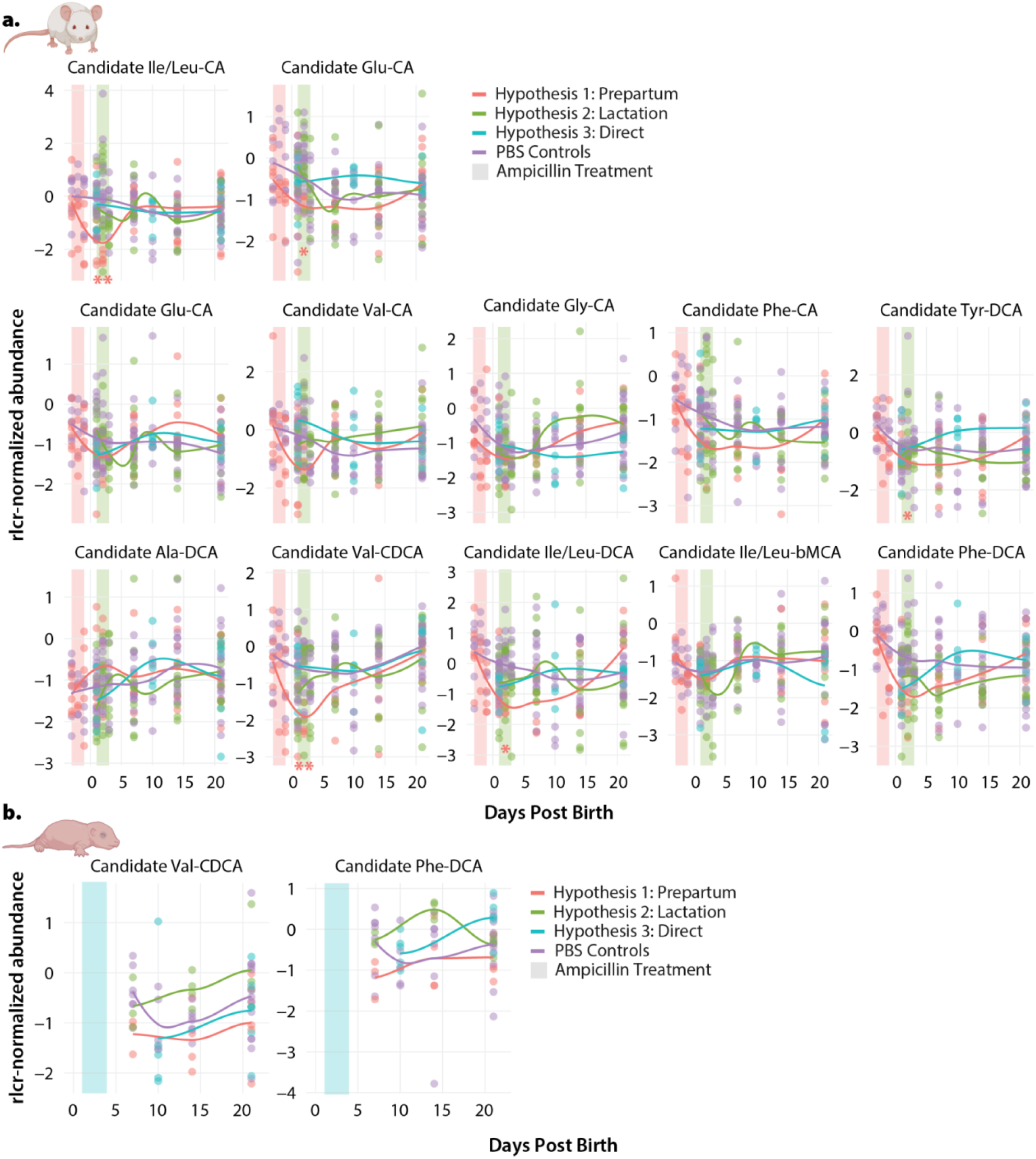
Abundance of candidate amino-acid conjugated bile acids in **a**. parent and **b**. infant mouse feces. Lines indicate average abundance across the entire sample group, points indicate abundance of individual samples. Abundance is shown as robust centered log-transformed (rlcr-transformed) values. Only molecules detected in at least 20% of samples are shown. Significance calculated using Tukey’s HSD with BH correction. p.adj < 0.05 = *, p.adj < 0.01 = **, p.adj < 0.001 = ***.

**Figure S5:**
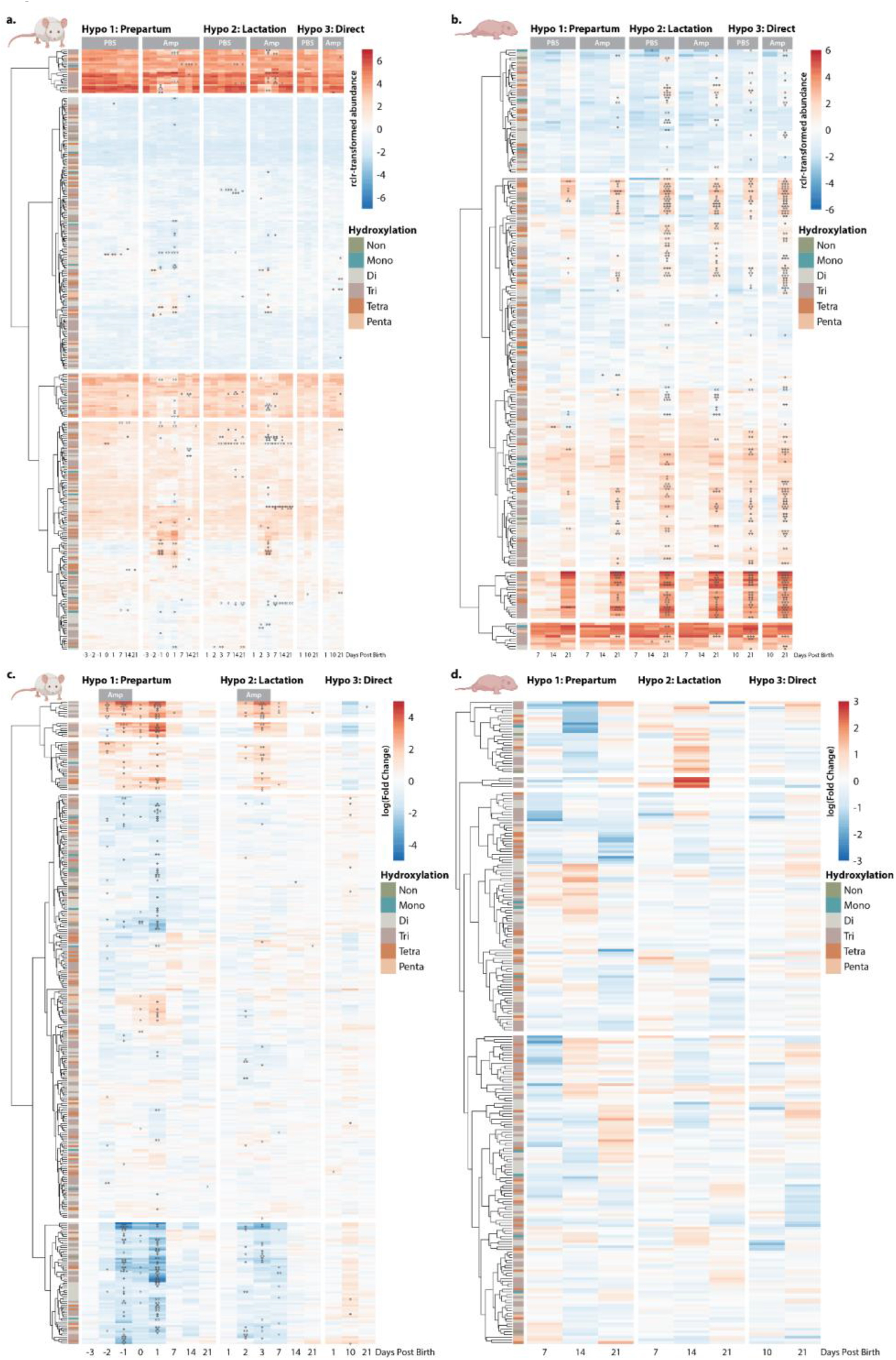
Heatmap of all candidate bile acids over time. **a-b**. Average abundance of bile acids in **a**. parent and **b**. infant feces. For **a-b** * indicates significance compared to first time point. **c-d**. Fold changes of bile acids in **a**. parent and **b**. infant feces. For **c-d** * indicates significance compared to PBS controls at the same time point. Significance calculated using Tukey’s HSD with BH correction. p.adj < 0.05 = *, p.adj < 0.01 = **, p.adj < 0.001 = ***. Annotations and individual p values are listed in **Table S1**.

**Figure S6:**
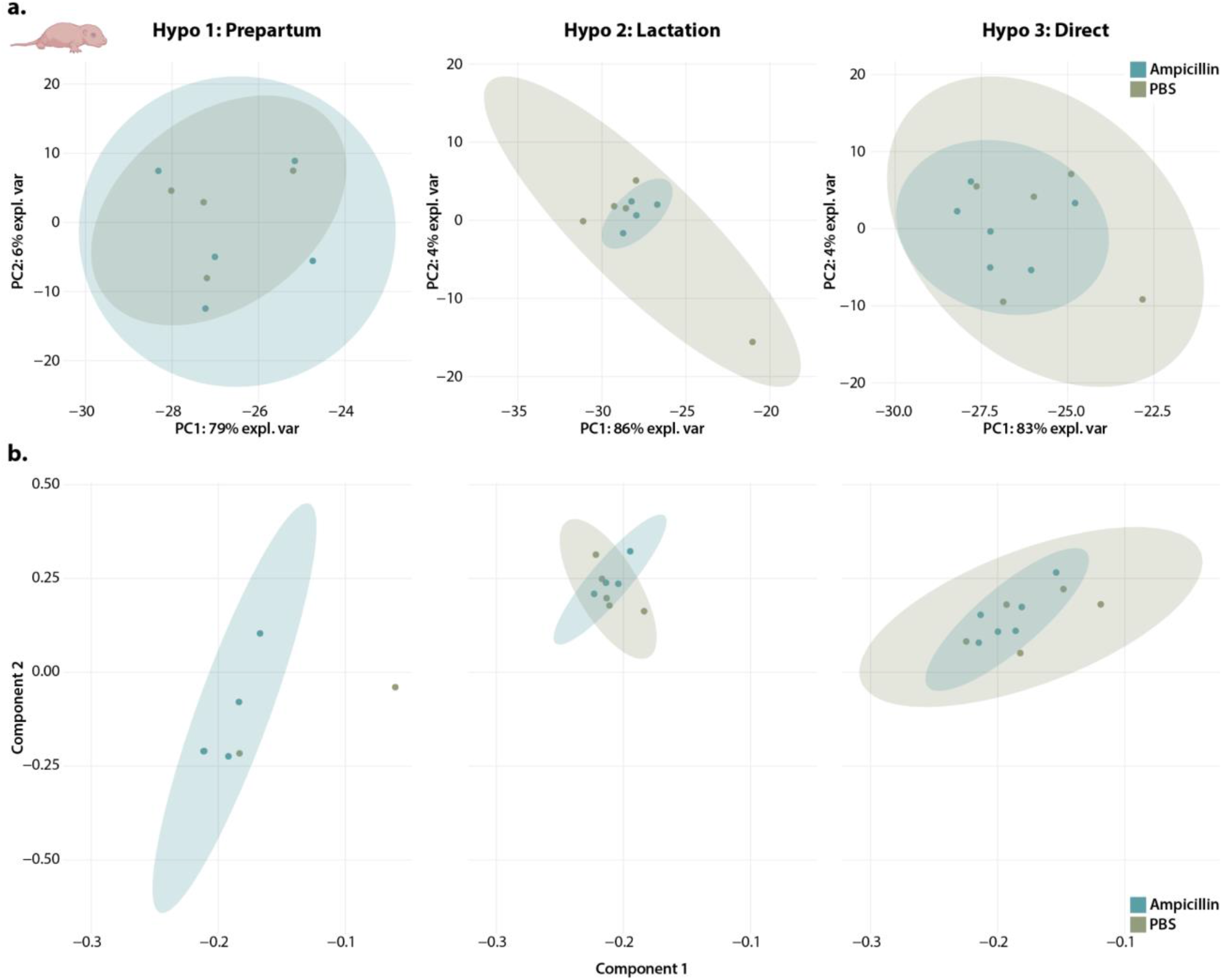
Principal components analysis (PCA) does not separate infant mice exposed to ampicillin. **a**. Standard PCA plot of infant mice treated with ampicillin or PBS controls **b**. Principal components from TEMPoral TEnsor Decomposition (TEMPTED), a longitudinal principal components analysis, using data from all timepoints. (Note, only two subjects in the Hypothesis 1 control groups had enough data to perform TEMPTED, thus no ellipse was drawn for this group).

## References

1. Naidoo, S., Bangalee, V. & Oosthuizen, F. Antibiotic use amongst pregnant women in a public hospital in KwaZulu-Natal. Health SA 26, 1516 (2021).

2. Andrade, S. E. et al. Prescription drug use in pregnancy. American Journal of Obstetrics and Gynecology 191, 398–407 (2004).

3. Baraka, M. A. et al. Patterns of infections and antimicrobial drugs’ prescribing among pregnant women in Saudi Arabia: a cross sectional study. J Pharm Policy Pract 14, 9 (2021).

4. Engeland, A. et al. Trends in prescription drug use during pregnancy and postpartum in Norway, 2005 to 2015. Pharmacoepidemiology and Drug Safety 27, 995–1004 (2018).

5. Prusakov, P. et al. A global point prevalence survey of antimicrobial use in neonatal intensive care units: The no-more-antibiotics and resistance (NO-MAS-R) study. EClinicalMedicine 32, 100727 (2021).

6. Schulman, J. et al. Newborn Antibiotic Exposures and Association With Proven Bloodstream Infection. Pediatrics 144, e20191105 (2019).

7. Mustafa, Z. U., Khan, A. H., Salman, M., Syed Sulaiman, S. A. & Godman, B. Antimicrobial Utilization among Neonates and Children: A Multicenter Point Prevalence Study from Leading Children’s Hospitals in Punjab, Pakistan. Antibiotics (Basel) 11, 1056 (2022).

8. Rooks, M. G. & Garrett, W. S. Gut microbiota, metabolites and host immunity. Nat Rev Immunol 16, 341–352 (2016).

9. Durack, J. & Lynch, S. V. The gut microbiome: Relationships with disease and opportunities for therapy. J Exp Med 216, 20–40 (2019).

10. Schwartz, D. J., Langdon, A. E. & Dantas, G. Understanding the impact of antibiotic perturbation on the human microbiome. Genome Medicine 12, 82 (2020).

11. Yao, Y. et al. The Role of Microbiota in Infant Health: From Early Life to Adulthood. Frontiers in Immunology 12, (2021).

12. Gensollen, T., Iyer, S. S., Kasper, D. L. & Blumberg, R. S. How colonization by microbiota in early life shapes the immune system. Science 352, 539–544 (2016).

13. Stewart, C. J. et al. Temporal development of the gut microbiome in early childhood from the TEDDY study. Nature 562, 583–588 (2018).

14. Yassour, M. et al. Natural history of the infant gut microbiome and impact of antibiotic treatment on bacterial strain diversity and stability. Sci Transl Med 8, 343ra81 (2016).

15. Bokulich, N. A. et al. Antibiotics, birth mode, and diet shape microbiome maturation during early life. Science Translational Medicine 8, 343ra82-343ra82 (2016).

16. Greenwood, C. et al. Early empiric antibiotic use in preterm infants is associated with lower bacterial diversity and higher relative abundance of Enterobacter. J Pediatr 165, 23–29 (2014).

17. Cox, L. M. et al. Altering the intestinal microbiota during a critical developmental window has lasting metabolic consequences. Cell 158, 705–721 (2014).

18. Roubaud-Baudron, C. et al. Long-Term Effects of Early-Life Antibiotic Exposure on Resistance to Subsequent Bacterial Infection. mBio 10, e02820–19 (2019).

19. Vrbanac, A. et al. Evaluating Organism-Wide Changes in the Metabolome and Microbiome following a Single Dose of Antibiotic. mSystems 5, e00340–20 (2020).

20. Wilkins, A. T. & Reimer, R. A. Obesity, Early Life Gut Microbiota, and Antibiotics. Microorganisms 9, 413 (2021).

21. Trasande, L. et al. Infant antibiotic exposures and early-life body mass. Int J Obes 37, 16–23 (2013).

22. Chen, R.-A. et al. Dietary Exposure to Antibiotic Residues Facilitates Metabolic Disorder by Altering the Gut Microbiota and Bile Acid Composition. mSystems 7, e00172–22 (2022).

23. Vatanen, T. et al. The human gut microbiome in early-onset type 1 diabetes from the TEDDY study. Nature 562, 589–594 (2018).

24. Jawad, A. B., Jansson, S., Wewer, V. & Malham, M. Early Life Oral Antibiotics Are Associated With Pediatric-Onset Inflammatory Bowel Disease-A Nationwide Study. J Pediatr Gastroenterol Nutr 77, 366–372 (2023).

25. Mitre, E. et al. Association Between Use of Acid-Suppressive Medications and Antibiotics During Infancy and Allergic Diseases in Early Childhood. JAMA Pediatrics 172, e180315 (2018).

26. Tamburini, S., Shen, N., Wu, H. C. & Clemente, J. C. The microbiome in early life: implications for health outcomes. Nat Med 22, 713–722 (2016).

27. Shekhar, S. & Petersen, F. C. The Dark Side of Antibiotics: Adverse Effects on the Infant Immune Defense Against Infection. Front Pediatr 8, 544460 (2020).

28. Korpela, K. et al. Intestinal microbiome is related to lifetime antibiotic use in Finnish pre-school children. Nat Commun 7, 10410 (2016).

29. Uzan-Yulzari, A. et al. Neonatal antibiotic exposure impairs child growth during the first six years of life by perturbing intestinal microbial colonization. Nat Commun 12, 443 (2021).

30. Quinn, R. A. et al. Global chemical effects of the microbiome include new bile-acid conjugations. Nature 579, 123–129 (2020).

31. Gentry, E. C. et al. Reverse metabolomics for the discovery of chemical structures from humans. Nature 1–3 (2023).doi:10.1038/s41586-023-06906-8

32. Patterson, A. et al. Bile Acids Are Substrates for Amine N-Acyl Transferase Activity by Bile Salt Hydrolase. (2022).doi:10.21203/rs.3.rs-2050120/v1

33. Mohanty, I. et al. The Underappreciated Diversity of Bile Acid Modifications. (2023).doi:10.2139/ssrn.4436846

34. Fu, T. et al. Paired microbiome and metabolome analyses associate bile acid changes with colorectal cancer progression. Cell Rep 42, 112997 (2023).

35. Prevention of Group B Streptococcal Early-Onset Disease in Newborns: ACOG Committee Opinion, Number 797. Obstet Gynecol 135, e51–e72 (2020).

36. Dalton, E. & Castillo, E. Post partum infections: A review for the non-OBGYN. Obstet Med 7, 98–102 (2014).

37. World Health Organization Pocket Book of Hospital Care for Children: Guidelines for the Management of Common Childhood Illnesses. (World Health Organization, Geneva, 2013).

38. Shi, P. et al. Time-Informed Dimensionality Reduction for Longitudinal Microbiome Studies. 2023.07.26.550749 (2023).doi:10.1101/2023.07.26.550749

39. Chiang, J. Y. L. Bile Acid Metabolism and Signaling. Compr Physiol 3, 1191–1212 (2013).

40. Thangamani, S. et al. Bile Acid Regulates the Colonization and Dissemination of Candida albicans from the Gastrointestinal Tract by Controlling Host Defense System and Microbiota. Journal of Fungi 7, 1030 (2021).

41. Guinan, J. & Thangamani, S. Antibiotic-induced alterations in taurocholic acid levels promote gastrointestinal colonization of Candida albicans. FEMS Microbiology Letters 365, fny196 (2018).

42. Devkota, S. et al. Dietary-fat-induced taurocholic acid promotes pathobiont expansion and colitis in Il10-/-mice. Nature 487, 104–108 (2012).

43. Wu, T. et al. Effects of Taurocholic Acid on Glycemic, Glucagon-like Peptide-1, and Insulin Responses to Small Intestinal Glucose Infusion in Healthy Humans. The Journal of Clinical Endocrinology & Metabolism 98, E718–E722 (2013).

44. Liu, Z. et al. Taurocholic acid is an active promoting factor, not just a biomarker of progression of liver cirrhosis: evidence from a human metabolomic study and in vitro experiments. BMC Gastroenterology 18, 112 (2018).

45. Yang, Z. et al. Serum Metabolomic Profiling Identifies Key Metabolic Signatures Associated With Pathogenesis of Alcoholic Liver Disease in Humans. Hepatology Communications 3, 542

46. Ovadia, C. et al. Association of adverse perinatal outcomes of intrahepatic cholestasis of pregnancy with biochemical markers: results of aggregate and individual patient data meta-analyses. The Lancet 393, 899–909 (2019).

47. Chen, Y., Li, H., Guo, H. & Zhou, J. Serum Taurocholic Acid Levels Have Predictive Value for Adverse Maternal and Infant Outcomes in Pregnant Women with Intrahepatic Cholestasis of Pregnancy: A Prospective Cohort Study. CEOG 50, 257 (2023).

48. Yang, Z. et al. Application of metabolomics in intrahepatic cholestasis of pregnancy: a systematic review. European Journal of Medical Research 27, 178 (2022).

49. Mertens, K. L., Kalsbeek, A., Soeters, M. R. & Eggink, H. M. Bile Acid Signaling Pathways from the Enterohepatic Circulation to the Central Nervous System. Frontiers in Neuroscience 11, (2017).

50. Li, M. et al. Gut microbiota-bile acid crosstalk contributes to the rebound weight gain after calorie restriction in mice. Nat Commun 13, 2060 (2022).

51. Wu, W. et al. Akkermansia muciniphila alleviates high-fat-diet-related metabolic-associated fatty liver disease by modulating gut microbiota and bile acids. Microbial Biotechnology 16, 1924–1939 (2023).

52. Bonde, Y., Eggertsen, G. & Rudling, M. Mice Abundant in Muricholic Bile Acids Show Resistance to Dietary Induced Steatosis, Weight Gain, and to Impaired Glucose Metabolism. PLOS ONE 11, e0147772 (2016).

53. Million, M. et al. Lactobacillus reuteri and Escherichia coli in the human gut microbiota may predict weight gain associated with vancomycin treatment. Nutr & Diabetes 3, e87–e87 (2013).

54. Nothias, L.-F. et al. Feature-based molecular networking in the GNPS analysis environment. Nat Methods 17, 905–908 (2020).

55. Schmid, R. et al. Integrative analysis of multimodal mass spectrometry data in MZmine 3. Nat Biotechnol 41, 447–449 (2023).

